# Microtubule stability modulates Schlemm’s canal cell mechanobiology and outflow facility in glaucoma

**DOI:** 10.64898/2026.04.27.721135

**Authors:** Haiyan Li, Nina Sara Fraticelli-Guzmán, Kristin M. Perkumas, Micah Chrenek, Andrew J. Feola, W. Daniel Stamer, C. Ross Ethier

**Author notes:** To whom correspondence should be addressed: C. Ross Ethier, PhD, Professor.

## Abstract

**Purpose:** The inner wall of Schlemm’s canal (SC) is a mechanosensitive endothelial monolayer that provides resistance to conventional aqueous humor drainage, a process dependent on pore formation. This study examined how microtubule (MT) stability affects SC cell mechanobiology, transcellular pore formation, and aqueous humor outflow dynamics.

**Methods:** MT stability in cultured SC cells from normal and glaucomatous human donors was manipulated pharmacologically. Changes in MT acetylation, phosphorylated myosin light chain, and F-actin were assessed by immunofluorescence and immunoblotting. GEF-H1 was knocked down using siRNA. Cellular stiffness was measured by atomic force microscopy. Transcellular pore formation was quantified using an established pore formation assay. Outflow facility was measured in enucleated mouse eyes using the iPerfusion system.

**Results:** MT stabilization in normal SC cells decreased actomyosin contractility and cellular stiffness, whereas MT destabilization increased contractility and stiffness; these effects involved the MT-associated Rho guanine nucleotide exchange factor GEF-H1. MT stability was also mechano-responsive to substrate stiffness. Furthermore, SC cells derived from glaucomatous donors exhibited reduced MT stability compared with normal SC cells. MT stabilization increased transcellular pore formation in both normal and glaucomatous SC cells. In *ex vivo* mouse eyes, paclitaxel perfusion to stabilize MTs significantly increased outflow facility relative to contralateral control eyes.

**Conclusions:** Our data suggest that MT stability influences SC cell contractility, stiffness, and transcellular pore formation and can alter aqueous humor outflow facility. These findings identify MT-dependent cytoskeletal remodeling as an important contributor to the biomechanics of the conventional outflow pathway and suggest that MT-associated pathways may represent potential targets for improving outflow function in glaucoma.

## Introduction

Glaucoma, a leading cause of irreversible blindness worldwide, is a progressive, age-related optic neuropathy^1^. Elevated intraocular pressure (IOP) is a primary, and currently the only modifiable, risk factor for glaucoma progression. The trabecular meshwork (TM) and Schlemm’s canal (SC) endothelium together comprise the central functional unit of the conventional aqueous humor (AH) outflow pathway, which provides resistance to AH drainage and therefore plays a key role in determining IOP^2,3^. SC is an endothelial-lined vessel whose cells display phenotypic characteristics of both vascular and lymphatic endothelia^4–8^. The SC inner wall endothelium facilitates AH outflow by forming openings in its monolayer which are of two distinct types: paracellular pores, which open at cell-cell junctions, and transcellular pores, which typically develop through giant vacuole-associated transcellular channels within individual SC endothelial cells. More specifically, the basal-to-apical pressure gradient between the anterior chamber and the SC lumen deforms SC endothelial cells to generate giant vacuoles and transcellular pores^7,9^. In glaucoma, decreases in SC inner wall pore size and density have been consistently observed^10,11^, implicating impaired pore formation as a potential contributor to elevated outflow resistance and ocular hypertension.

To assess transcellular pore formation in SC cells, our laboratory previously developed an *in vitro* assay that applies localized basal-to-apical cellular stretching using microparticles, thereby recapitulating the key *in vivo* mechanical features of giant vacuole formation^12^. Using this pore formation and detection assay, we demonstrated that basal cellular stretch imposed by particles underlying SC cells induces transcellular pore formation, and that treatments known to modulate intraocular pressure or outflow facility correspondingly alter transcellular pore formation. Moreover, pore formation was found to be closely associated with cellular stiffness, actomyosin contractility, and intracellular Ca^2+^ levels^12,13^.

The cytoskeleton, composed of actin filaments, microtubules (MTs), and intermediate filaments, is the principal determinant of cellular stiffness. In SC cells, actin polymerization is a key modulator of stiffness, with actin depolymerization reducing stiffness, whereas polymerization enhances it^13–15^. In contrast, the contribution of MTs to SC cell biomechanics remains largely unexplored in the context of glaucoma pathology. MTs are distinguished by their rapid dynamic reorganization^16–18^, enabling them to respond to intracellular and extracellular cues. Beyond their well-established roles in maintaining cell architecture^19^, vesicular transport^20^, and cell migration^21,22^, MTs function as mechano-active elements that integrate and transduce mechanical forces to mechanosensitive signaling pathways^23,24^. Importantly, MT stability is a critical determinant of this mechanobiological function, influencing cell contractility^24–26^ and stiffness^23,27^, both of which are closely linked to transcellular pore formation in SC cells and AH outflow. Thus, MT mechanobiology, and specifically MT stability, represents a critical yet understudied modulator of SC cell function and outflow dynamics.

Here, we investigated the role of MT stability in SC cells, with a particular focus on its modulation of actomyosin contractility, cellular stiffness, and mechanotransduction in response to substrate stiffness. Functionally, we examined how MT stabilization affects transcellular pore formation using our well-established *in vitro* assay, and how these mechanobiological processes ultimately influence AH outflow facility.

## Methods

### SC cell isolation and culture

Primary human SC cells were isolated from ostensibly normal and glaucomatous human donor eyes, as previously described, and were cultured according to established protocols ^8,28^. A total of five normal SC cell (nSC) strains isolated from healthy donor eyes and four glaucomatous cell (gSC) strains isolated from donors with a history of glaucoma were used in this study (**Table 1**). All SC cell strains were characterized based upon their typical spindle-like elongated cell morphology, expression of vascular endothelial-cadherin and fibulin-2, a net transendothelial electrical resistance of 10 ohms·cm^2^ or greater, and lack of myocilin induction following exposure to dexamethasone. Different combinations of SC cell strains were used per experiment depending on cell availability, and all studies were conducted between cell passages 3-6. SC cells were cultured in low-glucose Dulbecco’s Modified Eagle’s Medium (DMEM; Gibco; Thermo Fisher Scientific) containing 10% fetal bovine serum (FBS; HyClone, Thermo Fisher Scientific) and 1% penicillin/streptomycin/glutamine (PSG; Gibco) and maintained at 37°C in a humidified atmosphere with 5% CO_2_. Fresh media was supplied every 2-3 days.

**Table 1.** SC cell strain information. NM = not measured. N/A = not applicable. The first letter of the ID indicates normal (“n”) or glaucomatous (“g”). Outflow facility was measured in postmortem donor eyes.

| ID | Donor Age | Donor Sex | Facility<br>(μL/min/<br>mm Hg) | Ophthalmic<br>Medications |
| --- | --- | --- | --- | --- |
| nSC74 | 8 months | Male | NM | N/A |
| nSC75 | 10 years | Male | NM | N/A |
| nSC82 | 56 years | Male | NM | N/A |
| nSC87 | 62 years | Male | NM | N/A |
| nSC89 | 68 years | Male | NM | N/A |
| gSC57 | 78 years | Male | NM | Unknown |
| gSC64 | 78 years | Male | 0.07 | Unknown |
| gSC90 | 71 years | Female | 0.25 | Cosopt and<br>Latanoprost |
| gSC95 | 78 years | Male | 0.1 | N/A |

### Hydrogel preparation

To prepare soft hydrogel substrates, the hydrogel precursor gelatin methacryloyl (GelMA [6% w/v final concentration], Advanced BioMatrix, Carlsbad, CA, USA) was mixed with lithium phenyl-2,4,6-trimethylbenzoylphosphinate (LAP, 0.075% w/v final concentration) photoinitiator (Sigma-Aldrich, Saint Louis, MO). Stiff substrates also incorporated methacrylate-conjugated hyaluronic acid (MA-HA; [0.25% w/v final concentration], Advanced BioMatrix). Thirty microliters of the hydrogel solution were pipetted onto Surfasil-coated (Thermo Fisher Scientific) 18 × 18-mm square glass coverslips followed by placing 12-mm round silanized glass coverslips on top to facilitate even spreading of the polymer solution ^28–32^. Hydrogels were crosslinked by exposure to UV light (CL-3000 UV Crosslinker; Analytik Jena, Germany) at 1J/cm^2^. The hydrogel thickness was approximately 265 µm, calculated as the solution volume divided by the surface area of the gel on the coverslip. The hydrogel-adhered coverslips were removed with fine-tipped tweezers and placed hydrogel-side facing up in 24-well culture plates (Corning; Thermo Fisher Scientific). All the hydrogel substrates were coated with fibronectin (5 μg/cm^2^; Advanced BioMatrix) at room temperature for 1 hour^13,33^.

### SC cell seeding and treatments

SC cells were seeded at 2 × 10^4^ cells/cm^2^ on premade hydrogel substrates, glass coverslips, or tissue culture plates and cultured in DMEM with 10% FBS and 1% PSG for one or two days until ∼80-90% confluence was reached. SC cell on coverslips or tissue culture plates were then exposed to reduced-serum media (DMEM with 1% FBS and 1% PSG) for 24 hours, followed by treatment with the indicated agents, including the MT stabilizer paclitaxel (10 µM; Sigma) or the MT destabilizer nocodazole (10 µM; Sigma). Cells cultured on hydrogels were similarly switched to DMEM with 1% FBS and 1% PSG for 2 days, followed by analysis of tubulin acetylation levels.

### Immunocytochemistry staining analysis

SC cells were fixed with 4% paraformaldehyde (PFA; Thermo Fisher Scientific) at room temperature for 15 minutes, permeabilized with 0.1% Triton™ X-100 for 15 minutes (Thermo Fisher Scientific), blocked with goat serum (BioGeneX, Fremont, CA, USA), and incubated with primary antibodies [anti-alpha Tubulin (acetyl K40), ab179484, Abcam, 1:1000; anti-alpha Tubulin, ab7291, Abcam, 1:1000; anti-phospho-myosin light chain 2 (Thr18/Ser19), 3764S, Cell Signaling Technology)] followed by incubation with fluorescent secondary antibody (Alexa Fluor® 488 anti-Rabbit, A27034, Invitrogen, 1:500; Alexa Fluor® 546 anti-Mouse, A11003, Invitrogen, 1:500). Nuclei were counterstained with 4′,6′-diamidino-2-phenylindole (5 μM, DAPI; Abcam, Waltham, MA, USA). Similarly, cells were stained with Phalloidin-iFluor 488 (Invitrogen; Thermo Fisher Scientific) or 594 (1:500, Cell signaling technology)/DAPI (Abcam) according to the manufacturer’s instructions. Glass coverslips/hydrogel substrates were mounted with ProLong™ Gold Antifade (Invitrogen) on Superfrost™ microscope slides (Fisher Scientific), and fluorescent images were acquired with a Leica DM6 B upright microscope system or with a Zeiss LSM 700 confocal microscope system.

All fluorescent image analyses were performed using FIJI software ^34^ (National Institutes of Health (NIH), Bethesda, MD, USA). Fluorescence intensity of all F-actin or phospho-myosin light chain 2 (Thr18/Ser19) within the image was measured with image background subtraction. We then normalized the intensity by cell number and calculated fold-difference vs. control cells.

### siRNA transfection

For siRNA-mediated GEFH1 depletion experiments, SC cells were seeded at 2 × 10^4^ cells/cm^2^ on glass coverslips, or tissue culture plates and cultured in DMEM with 10% FBS and 1% PSG. The following day, the cell culture medium was changed to antibiotic-free and serum-free DMEM and the samples were kept in culture for 24 h followed by transfection. Transfection was performed using a final concentration of 3% (v/v) lipofectamine RNAimax (Invitrogen; Thermo Fisher Scientific) with 150 nM RNAi (ON-TARGETplus non-targeting siRNA or ON-TARGETplus Human ARHGEF2 siRNA SMARTpool, Horizon/Dharmacon, Lafayette, CO, USA), according to the manufacturer’s instructions.

### Quantitative reverse transcription-polymerase chain reaction (qRT-PCR) analysis

Total RNA was extracted from SC cells using PureLink RNA Mini Kit (Invitrogen; Thermo Fisher Scientific). RNA concentration was determined with a NanoDrop spectrophotometer (Thermo Fisher Scientific). RNA was reverse transcribed using iScript™ cDNA Synthesis Kit (BioRad, Hercules, CA, USA). One hundred nanograms of cDNA were amplified in duplicates in each 40-cycle reaction using a CFX 96 Real Time PCR System (BioRad) with annealing temperature set at 60°C, Power SYBR™ Green PCR Master Mix (Thermo Fisher Scientific), and custom-designed qRT-PCR primers for GEFH1 (Forward: TTTCAGGCATGACCATGTGC; Reverse: TTGCTTCTGCTTGACCTTGG). Transcript levels were normalized to GAPDH, and mRNA fold-change calculated relative to mean control values using the comparative CT method.

### Immunoblot analysis

Protein was extracted from SC cells using lysis buffer (CelLytic^TM^ M, Sigma-Aldrich) supplemented with Halt™ protease/phosphatase inhibitor cocktail (Thermo Fisher Scientific). Equal protein amounts (15 µg), determined by standard bicinchoninic acid assay (Pierce; Thermo Fisher Scientific), in 4× loading buffer (Biorad) with 5% beta-mercaptoethanol (Fisher Scientific) were boiled for 5 min and subjected to SDS-PAGE using 4-15% Mini-PROTEAN® TGX™ gels (Biorad) at 200V for 30 min and transferred to 0.45 µm PVDF membranes (Sigma; Thermo Fisher Scientific). Membranes were blocked with blocking buffer (LI-COR Biosciences, Lincoln, NE) in tris-buffered saline with 0.2% Tween®20 (Thermo Fisher Scientific) and probed with primary antibodies followed by incubation with fluorescence-conjugated secondary antibodies. Bound antibodies were visualized with the imager (LI-COR Odyssey CLx and Biorad). Densitometry was performed using image studio and image lab software; data were normalized to total α tubulin followed by calculation of relative change vs. control.

### Cell and hydrogel stiffness measurement

SC cells were seeded at 2 × 10^4^ cells/cm^2^ on glass coverslips and cultured in DMEM with 10% FBS and 1% PSG for one or two days until cells reached ∼80-90% confluence. Then, SC cells were exposed to 10 μM paclitaxel or nocodazole for 30 minutes, as described above. An MFD-3D AFM (Asylum Research, Santa Barbara, CA, USA) was used to make stiffness measurements using silicon nitride cantilevers with an attached borosilicate sphere (diameter = 10 μm; nominal spring constant = 0.1 N/m; Novascan Technologies, Inc., Ames, IA, USA). Cantilevers were calibrated by measuring the thermally induced motion of the unloaded cantilever before measurements. The trigger force was set to 600 pN to avoid substrate effects and the tip velocity was adjusted to 800 nm/s to avoid viscous effects^35^. Five measurements/cell were conducted, and at least 5 cells were measured/group of one cell strain. For hydrogel stiffness measurement, a force map covering a 20 × 20 μm area (5 x 5 grid of points) was measured. Data from AFM measurements were fitted to the Hertz model to calculate the effective Young’s modulus of the cells, assuming the Poisson’s ratio was 0.5.

### Intracellular pore detection assay

16-well chambered glass bottom plates (Invitrogen) were plasma-treated and coated with 0.5 mg/mL biotinylated gelatin overnight at 37°C. The gelatin was then crosslinked with microbial transglutaminase (0.1 unit/µL, Sigma-Aldrich) at 37°C overnight. After UV sterilization of the plate, carboxyl ferromagnetic particles (4.0-4.9 µm; Spherotech Inc.) were added and incubated at room temperature for 10 minutes, followed by three gentle PBS washes. SC cells were seeded on top of the particles at a density of 7.5 × 10³ cells/cm² in DMEM supplemented with 10% FBS and 1% PSG, and incubated for 3.5 hours at 37°C with 5% CO_2_.

SC cells were treated with DMSO, 10 µM paclitaxel, or 10 µM nocodazole in DMEM supplemented with 10% FBS and 1% PSG for 30 minutes. Cells were then incubated with a fluorescent tracer (Streptavidin, Alexa Fluor™ 488 conjugate; Invitrogen) under the indicated treatment conditions (DMSO, 10 µM paclitaxel, or 10 µM nocodazole) in DMEM containing 10% FBS and 1% PSG for 5 minutes. The cells were then immediately washed three times with PBS, fixed with 4% formaldehyde for 15 minutes at room temperature, permeabilized with 0.1% Triton X for 15 minutes, and stained with Phalloidin-594 (Cell Signaling Technology) and DAPI. The glass cover was detached from the 16-well chambered system using a coverglass removal tool (Invitrogen), mounted on another glass slide with Prolong Gold Antifade Mountant (Invitrogen), and stored in the dark at 4°C until examination with a Leica DM6 B upright microscope system. Fluorescent and brightfield images were acquired using a 10x objective to generate tile scans for each well.

For the transcellular pore formation analysis, a 4 mm diameter circle was delineated at the center of each tile-scanned well image to exclude cells near the well edges, restricting the analysis to cells within this circle. The numbers of cells, particles beneath cells, and pores co-localized with particles were manually counted. The percentage of particle-induced pores was quantified as the number of pores associated with particles divided by the number of particles under cells, multiplied by 100^36^.

### Outflow facility measurement

Outflow facility, defined as the mathematical inverse of conventional outflow resistance, was measured in enucleated eyes using the previously established iPerfusion system^37^. Eight C57BL/6J mice (3 males and 5 females, 2-4 months old; Jackson Laboratory, Bar Harbor, ME, USA) were used in this study. Mice were euthanized by CO_2_ inhalation, after which eyes were carefully enucleated and mounted in an eye holder positioned at the center of the perfusion chamber. The temperature-controlled chamber (35°C) was filled with prewarmed Dulbecco’s phosphate-buffered saline supplemented with 5.5 mM D-glucose (DBG), fully submerging the eyes. System sensors were calibrated before each measurement session to ensure accuracy and to confirm the absence of air bubbles or leaks, which can introduce substantial measurement errors. Perfusion needles were filled with filtered DBG containing 3% (2-Hydroxypropyl)-β-cyclodextrin (Sigma) to enhance drug solubility and supplemented with either paclitaxel (100 µM) or vehicle control (1% DMSO).

The paclitaxel concentration of 100 μM in the perfusate was higher that the concentration used in *in vitro* experiments so as to account for drug dilution within the anterior chamber. More specifically, without a 2-needle anterior chamber exchange, the drug concentration delivered to the outflow pathway is invariably less than the nominal concentration. We estimated this dilution using calculations (Dr. Joseph M. Sherwood, Imperial College London; personal communication) based on an assumed anterior chamber volume of 3 μL and expected flow rates of 16 nL/min at 8 mmHg^37,38^ to estimate a paclitaxel concentration in the conventional outflow pathway of ∼20 μM after 45 minutes of preconditioning.

Following cannulation, one eye was perfused with paclitaxel, while the contralateral eye was perfused with DMSO as the vehicle control at 8 mmHg for 45 minutes (preconditioning period) before facility measurements were initiated. Outflow facility was then determined using a nine-step pressure protocol starting at 5 mmHg, increasing in 1.5 mmHg increments up to 17 mmHg, and ending with a final step at 8 mmHg. The resulting flow-pressure data were fit with an empirical power-law relationship [Q(P) = C_r_(P/P_r_)^β^], where P is the measured steady-state pressure, Q is the measured steady-state flow rate at each pressure step, and P_r_ is the reference pressure (8 mmHg), which approximates the physiological pressure drop across the conventional outflow pathway in living mice^37^. C_r_ is the facility evaluated at P_r_ = 8 mmHg, and β is a nonlinearity parameter determined when fitting the model to the data. Based on the facility data, we quantified the net effect of paclitaxel by computing 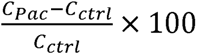, where *C* is reference facility and subscripts denote paclitaxel-treated (Pac) and control eyes (ctrl). This computed quantity can be interpreted as the relative difference in outflow facility between control and paclitaxel-treated eyes, expressed as a percentage.

### Immunohistochemistry analysis

Following *ex vivo* perfusion, eyes were fixed by immersion in 4% PFA overnight at 4°C. PFA was then removed, and eyes were washed with DPBS. Three pairs of eyes (both control and paclitaxel-treated) showing a significant outflow facility response to the drug (i.e., net facility change > 30%) were used for immunohistochemistry studies. Eyes were hemisected along the equator and the posterior segments and lenses removed. The anterior segments were then cut into four quadrants. Each quadrant was placed in 30% sucrose overnight at 4°C, embedded in Tissue-Plus^TM^ O.C.T. Compound (Fisher Scientific) in a 10 × 10 × 5-mm cryomold. Blocks were cut to create 10-µm sections using a cryostat (Leica Biosystems Inc., Buffalo Grove, IL, USA). Sections were placed on positively charged slides, followed by incubating in a 37°C oven for 10 minutes. Sections were then blocked with 10% goat serum (Gibco; Thermo Fisher Scientific) and incubated with primary antibodies [anti-CD31, MAB1398Z, Sigma-Aldrich, 1:100; anti-alpha Tubulin (acetyl K40), ab179484, Abcam, 1:200; anti-alpha Tubulin, ab7291, Abcam, 1:200], followed by incubation with fluorescent secondary antibody (Alexa Fluor® 488 anti-American Hamster, Invitrogen, 1:500; Alexa Fluor® 546 anti-Rabbit, Invitrogen, 1:500; Alexa Fluor® 647 anti-Mouse, Invitrogen, 1:500). Nuclei were counterstained with 4′,6′-diamidino-2-phenylindole (5 μM, DAPI; Abcam, Waltham, MA, USA). Slides were mounted with ProLong^TM^ Gold Antifade (Thermo Fisher Scientific), and fluorescent images were acquired with a Nikon Ti2 microscope with A1R confocal as z- stack with 1 μm step interval. The trabecular meshwork and SC inner wall region was outlined using merged brightfield, DAPI, and CD3 images. Fluorescence intensity of acetylated-α tubulin (ac-α tubulin) and total α tubulin was measured within the region of interest from maximum intensity projections using FIJI software (NIH) with image background subtraction, The Ac-α-tubulin/total α-tubulin ratio was then calculated and expressed as fold change vs. control.

### Statistical analysis

Statistical analysis was conducted using GraphPad Prism (v11.0.1; Boston, MA), with a significance (Type I error) threshold set at p = 0.05. Sample sizes are specified in each figure caption. All data sets, except for AFM and iPerfusion, were tested for normality using the Shapiro-Wilk test and were confirmed to meet the normality criteria (p>0.05). The AFM and iPerfusion data sets, which are expected to be log-normally distributed^37,39^, were tested for log-normality using the Shapiro-Wilk test on log-transformed data, and met the relevant criteria (p>0.05). Comparisons between groups were assessed by t-test, one-way or two-way analysis of variance (ANOVA) with Tukey’s multiple comparisons *post hoc* tests, or a suitable nonparametric test, as appropriate.

## Results

### MT stability affects SC cell actomyosin contractility

Schlemm’s canal (SC) cells are highly contractile, and their contractile state is closely associated with outflow facility and IOP^28,31^. To determine the role of MT stability, normal SC cells were treated with the MT stabilizer paclitaxel or the MT destabilizer nocodazole for 30 minutes. We confirmed that paclitaxel and nocodazole increased and decreased MT stability in SC cells, respectively (Fig. 1A, B; Suppl. Fig. 1), as assayed using MT acetylation as an established surrogate marker of MT stability^40,41^. To assess the impact of MT stability on SC cell contractility, we examined two key actomyosin-related markers: phosphorylated myosin light chain (p-MLC) levels and filamentous actin (F-actin) levels. Over short time scales (30 minutes), paclitaxel treatment significantly reduced p-MLC levels by 35.1% (p < 0.01; Fig. 1C, D) without affecting F-actin levels (Fig. 1E, F). In contrast, nocodazole treatment significantly increased both p-MLC (1.9-fold, p < 0.0001; Fig. 1C, D) and F-actin levels (1.9-fold, p < 0.0001; Fig. 1E, F), indicating enhanced actomyosin contractility following MT destabilization. Furthermore, prolonged paclitaxel treatment (24 hours) markedly reduced F-actin stress fibers by 58.1% (p < 0.0001) and decreased p-MLC levels by 82.9% (p < 0.0001; Suppl. Fig. 2), suggesting that sustained MT stabilization is required to induce significant remodeling of F-actin. Together, these results demonstrate that MT stability negatively modulates actomyosin tone in SC cells.

**Fig. 1.**
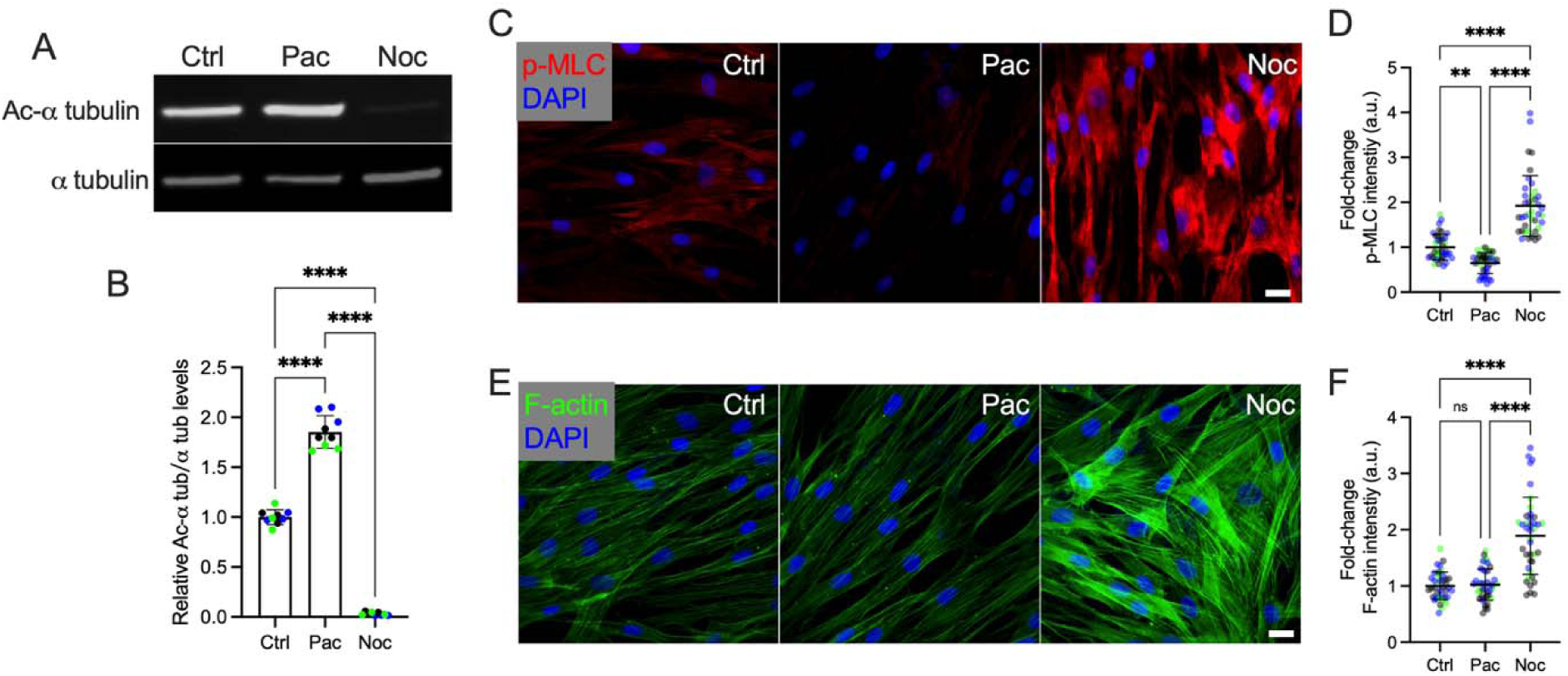
MT stability affects actomyosin contractility in normal Schlemm’s canal (nSC) cells. (A) Representative immunoblot showing acetylated-α tubulin (Ac-α tubulin) and total α tubulin in nSC cells treated for 30 minutes with control (DMSO), 10 μM paclitaxel (Pac; MT stabilizer), or 10 μM nocodazole (Noc; MT destabilizer). (B) Quantification of immunoblot expressed as normalized ac-α tubulin/α tubulin ratios (n = 9 replicates from 3 different nSC cell strains, with three independent replicates per strain). (C, E) Representative fluorescence micrographs of F-actin and phosphorylated myosin light chain (p-MLC) in nSC cells under the same treatment conditions for 30 minutes: control (DMSO), 10 μM paclitaxel (Pac), or 10 μM nocodazole (Noc). Scale bar, 20 μm. (D, F) Quantification of normalized F-actin and p-MLC fluorescence intensities (n = 30 images per group from 3 different nSC cell strains, with three independent replicates per strain). Symbols of the same color represent data from the same cell strain. Data are presented as mean ± SD. Statistical significance was determined using two-way ANOVA with multiple comparisons tests (**p < 0.01, ****p < 0.0001).

### MT stability mediates SC cell actomyosin contractility via GEF-H1

RhoA, a key member of the mammalian Rho GTPase family, is essential for actomyosin regulation. Upon activation, RhoA promotes the formation of actomyosin contractile structures through its downstream effector proteins, including formin, which drive linear actin polymerization, and Rho-associated kinase (ROCK), which phosphorylates the MLC to enhance myosin II–mediated contractility^42,43^. RhoA activation is mediated by guanine nucleotide exchange factors (GEFs)^44^, including GEF-H1, a MT-associated Rho GEF that remains inactive while bound to MTs but becomes activated and released upon MT destabilization^45,46^. To determine the possible role of GEF-H1 in MT-mediated SC cell contractility, we used siRNA to knock down GEF-H1 expression. Following 48 hours of siRNA treatment in nSC cells, GEF-H1 mRNA levels were reduced by 79% (Fig. 2A), with a corresponding 55.1% reduction in GEF-H1 protein levels (Fig. 2B).

**Fig. 2.**
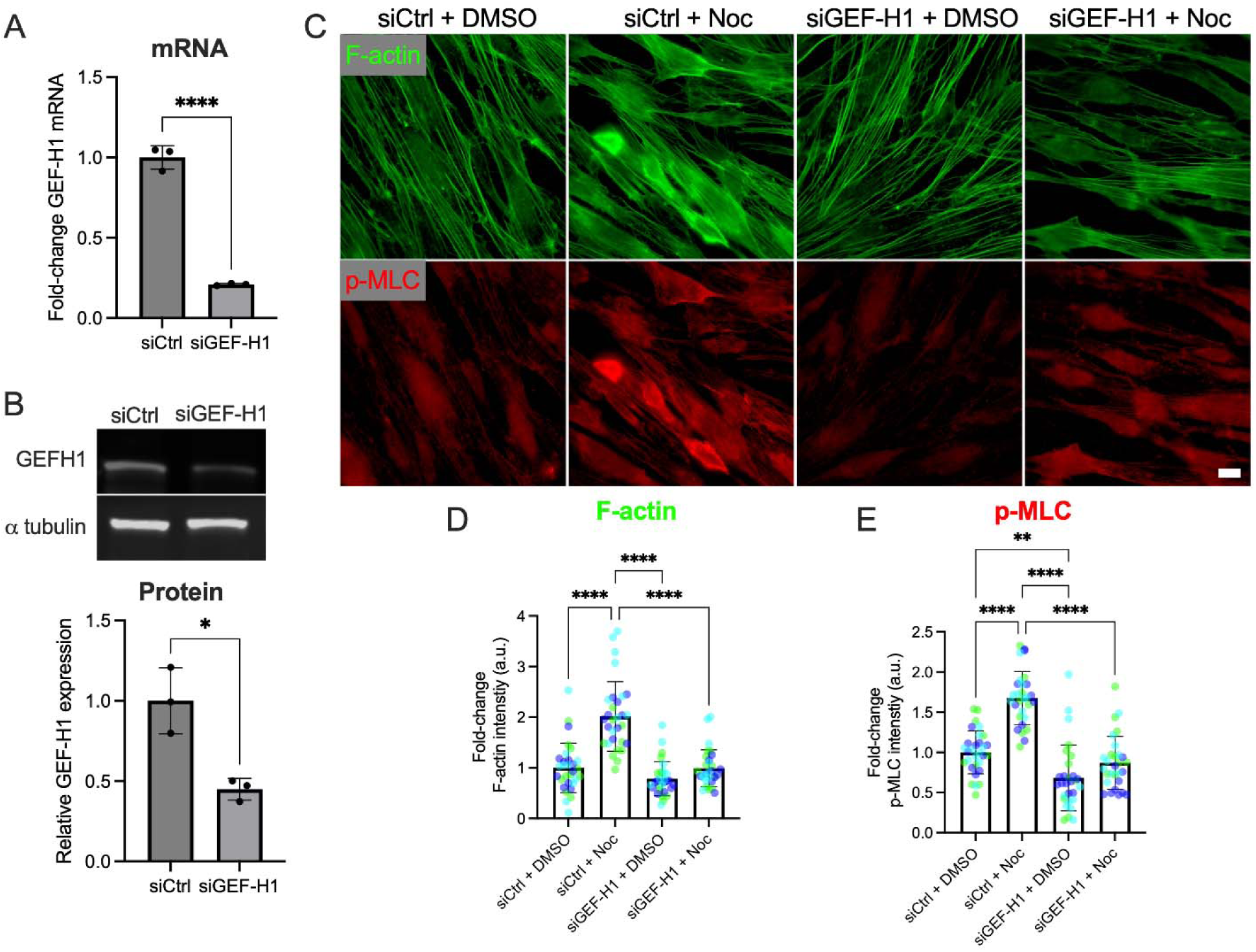
MT stability modulates actomyosin contractility in nSC cells via GEF-H1. (A) Normalized GEF-H1 mRNA levels in nSC cells are reduced following siGEF-H1 treatment for 48 hours, as determined by qRT-PCR (normalization to GAPDH levels followed by normalization to control siRNA treatment; n = 3 replicates from one cell strain). (B) Representative immunoblot (top) showing acetylated-α tubulin (ac-α tubulin) and total α tubulin in nSC cells, with quantification (bottom) expressed as normalized ac-α tubulin/α tubulin ratios following 48 hour control and GEF-H1 siRNA treatments (n = 3 replicates from one cell strain). (C) Representative fluorescence micrographs of F-actin and p-MLC in nSC cells under the following conditions: siControl (siCtrl) + DMSO, siCtrl + 10 μM nocodazole (Noc), siGEFH1 + DMSO, or siGEFH1 + 10 μM Noc. Cells were treated with siRNA for 48 hours followed by nocodazole treatment for 30 minutes. Scale bar, 20 μm. (D, E) Quantification of normalized F-actin and p-MLC fluorescence intensities (n = 30 images per group from 3 different nSC cell strains, with three independent replicates per strain). Symbols of the same color represent data from the same cell strain. Data are presented as mean ± SD. Statistical significance was determined using two-way ANOVA with multiple comparisons tests (**p < 0.01, ****p < 0.0001).

We observed that GEF-H1 knockdown significantly reduced p-MLC levels without altering F-actin levels (Fig. 2C-E). Consistent with our previous findings (Fig. 1), nocodazole treatment significantly increased both F-actin stress fibers and p-MLC levels; however, these effects were abolished by GEF-H1 knockdown. Notably, co-treatment with siGEF-H1 and nocodazole restored F-actin and p-MLC levels to those observed in siCtrl-treated cells under control conditions (Fig. 2C-E). Taken together, these findings identify GEF-H1 as a key mediator linking microtubule destabilization to RhoA–ROCK–dependent actomyosin contractility in SC cells.

### MT stability modulates SC cell stiffness and transcellular pore formation

Most AH crosses the continuous endothelial monolayer formed by SC inner wall cells by passing through micron-sized pores, and pore density is decreased in glaucoma^47^. Transcellular pore formation in SC cells has been shown to be related to cell stiffness and contractility^12,14,15^. We used atomic force microscopy (AFM) to measure normal SC cell stiffness, finding that MT stabilization decreased SC cell stiffness by 36.4% (p<0.01), whereas MT destabilization increased SC cell stiffness by 44.0% (p<0.001) (Fig. 3A). We then examined whether MT stability affects transcellular pore formation in normal SC cells. To assess transcellular pore formation, we used an *in vitro* cell-based assay previously developed by our lab, in which localized mechanical stretch is applied to SC cells^12,13^ by seeding cells on microparticles (4.0-4.9 μm in diameter). Transcellular pore formation, induced by this stretch, is detected using a fluorescently-tagged streptavidin which traverses SC cells at pore sites and irreversibly binds to an underlying biotinylated substrate, producing a persistent and detectable fluorescent signal. Consistent with our previous observations, normal SC cells formed transcellular pores in response to basal-to-apical stretch induced by microparticles (Fig. 3B). To investigate the role of MT stability in SC cell pore formation, cells were seeded onto particles and treated with either 10 µM paclitaxel (MT stabilizer) or 10 µM nocodazole (MT destabilizer) for 30 minutes, after which pore formation was assessed (Fig. 3C). Consistent with the changes in cell stiffness, MT stabilization increased transcellular pore formation rate from 9.0% to 11.5%, whereas MT destabilization reduced pore formation rate to 4.9% (Fig. 3D). Together, these results demonstrate that MT stability in SC cells plays a critical role in modulating cell stiffness and transcellular pore formation, suggesting that MT stability may influence outflow facility and IOP.

**Fig. 3.**
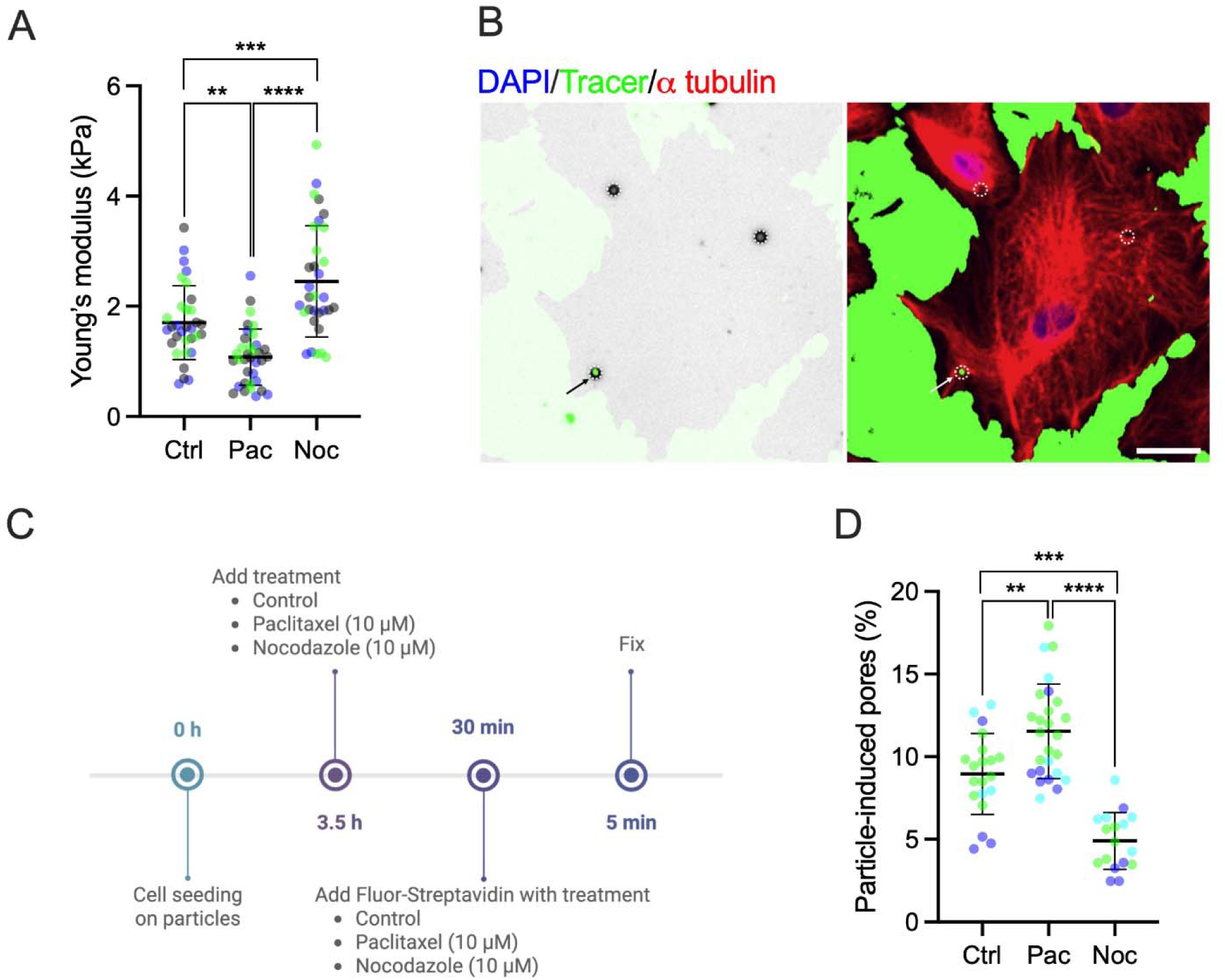
MT stability is associated with normal SC (nSC) cell stiffness and transcellular pore formation. (A) Young’s modulus measured by AFM in nSC cells exposed to control (DMSO), 10 μM paclitaxel (Pac), or 10 μM nocodazole (Noc) for 30 minutes (n = 32-35 cells per group from 3 different nSC cell strains). (B) The left panel shows a representative fluorescent micrograph merged with a brightfield image, allowing identification of microbead locations (white circles). The right panel shows the same field of view with the brightfield channel turned off, allowing identification of spots where fluorescently tagged streptavidin (green) has bound with the biotin substrate under cultured SC cells (at sites of intracellular pores) as well as surrounding regions not covered by cells. White circles outline particles. The arrow indicates a particle-induced pore. Scale bar: 20 μm. (C) Schematic showing the time course of the pore formation assay: SC cells were seeded atop particles on a biotinylated gelatin-coated glass substrate. Three and half hours after cell seeding, the relevant treatment (DMSO [control], 10 μM paclitaxel, or 10 μM Nocodazole) was introduced into the culture media and incubated for 30 mins. Then fluorescently-tagged streptavidin plus the relevant treatment were introduced into the culture media and incubated for 5 additional mins. Created with BioRender.com. (D) Pore incidence (ratio of pore-associated particles to particles under cells) in nSC cells (n = 17-26 wells from 3 different nSC cell strains). Symbols with different colors represent different cell strains. The lines and error bars indicate Mean ± SD. Significance was determined by two-way ANOVA using multiple comparisons (**p < 0.01, ***p < 0.001, ****p < 0.0001).

### MT stability is mechano-responsive to substrate stiffness in SC cells

The trabecular meshwork, which serves as the substrate for SC inner wall cells, has been shown to be significantly stiffer in glaucoma, with normal trabecular meshwork exhibiting a stiffness of approximately 0.5-9 kPa as measured by AFM, whereas glaucomatous tissue is approximately 1.5-20-fold stiffer depending on the measurement method and the location of measurement^48–51^. In our previous work, we developed a tissue-mimetic hydrogel system with defined elastic moduli of 2.36 kPa (soft) and 8.00 kPa (stiff), as measured by AFM^13^. These stiffness values were selected to model the physiological properties of normal trabecular meshwork tissue and the pathological stiffening observed in glaucoma. When normal SC cells were cultured on soft and stiff substrates, cells on the stiff substrate exhibited significantly lower levels of tubulin acetylation (30.4% decrease, p<0.0001) compared to those on the soft substrate, indicating reduced MT stability under glaucomatous-like stiff conditions (Fig. 4). Collectively, these findings suggest that MT stability in SC cells is mechano-responsive to substrate stiffness, and that targeting MT stability may represent a potential strategy to mitigate glaucoma-associated pathological changes.

**Fig. 4.**
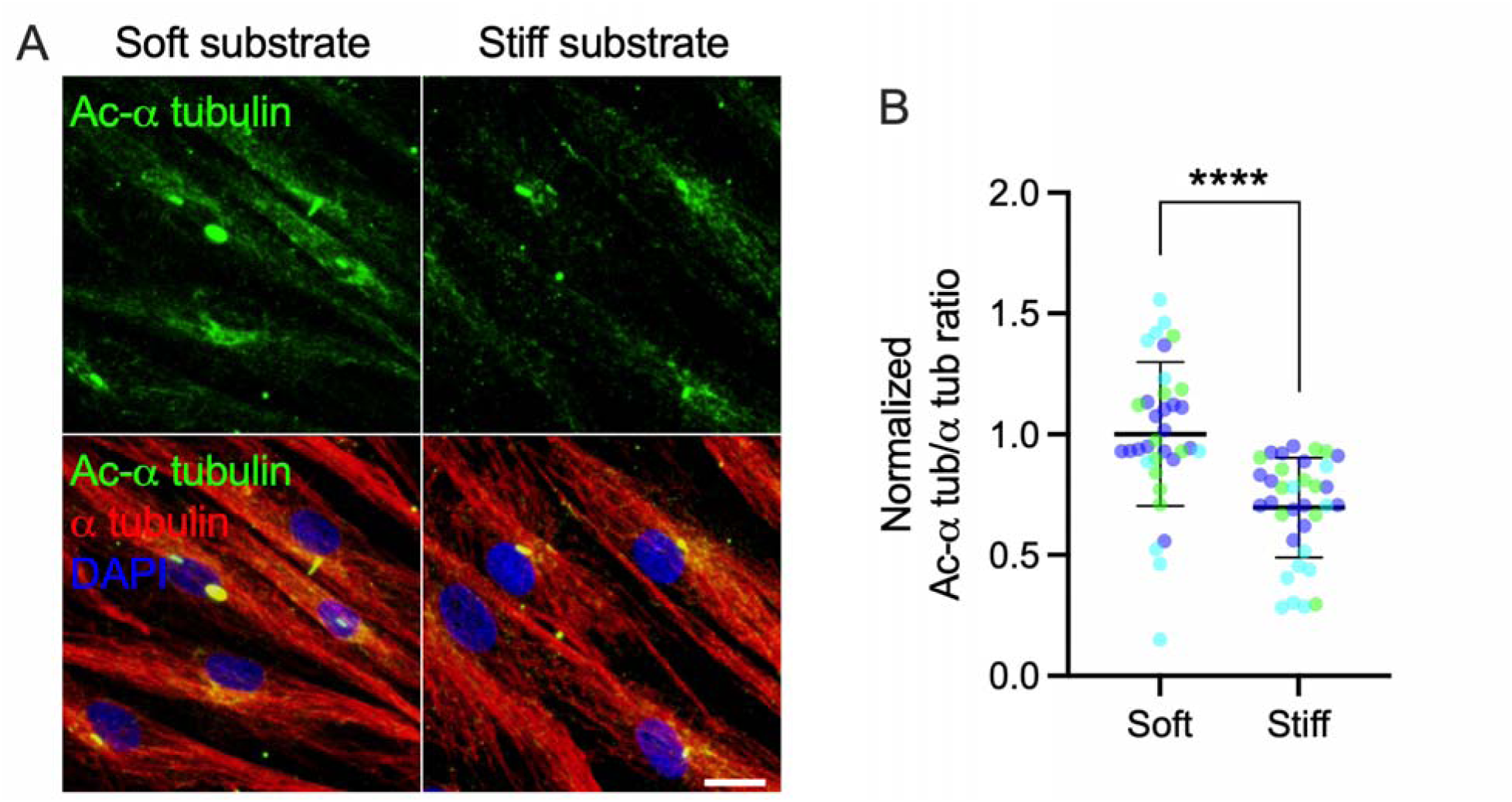
A glaucomatous-like stiff substrate reduces MT stability in normal SC (nSC) cells. (A) Representative fluorescence micrographs of acetylated-α tubulin (ac-α tubulin) and total α tubulin (α tubulin) in nSC cells on soft and stiff hydrogel substrates. Scale bar, 20 μm. (B) Quantification of normalized acetylated-α tubulin/total α tubulin fluorescence (n = 35 images per group from 3 different nSC cell strains, with three independent replicates per strain). Symbols of the same color represent data from the same cell strain. Data are presented as mean ± SD. Statistical significance was determined using two-way ANOVA with multiple comparisons tests (****p < 0.0001).

### Reduced MT stability in glaucomatous SC is restored by pharmacological stabilization

Glaucomatous SC (gSC) cells, derived from patients with glaucoma, are known to be stiffer and exhibit a more fibrotic phenotype compared to non-glaucomatous SC cells^13,15,52^. Here, we observed that gSC cells displayed lower levels of acetylated MTs compared to nSC cells across multiple cell strains (Fig. 5A, B, D, E). When data were pooled between normal strains and between glaucomatous strains, nSC cells exhibited significantly higher MT acetylation levels than gSC cells (p < 0.0001; Fig. 5C, F).

**Fig. 5.**
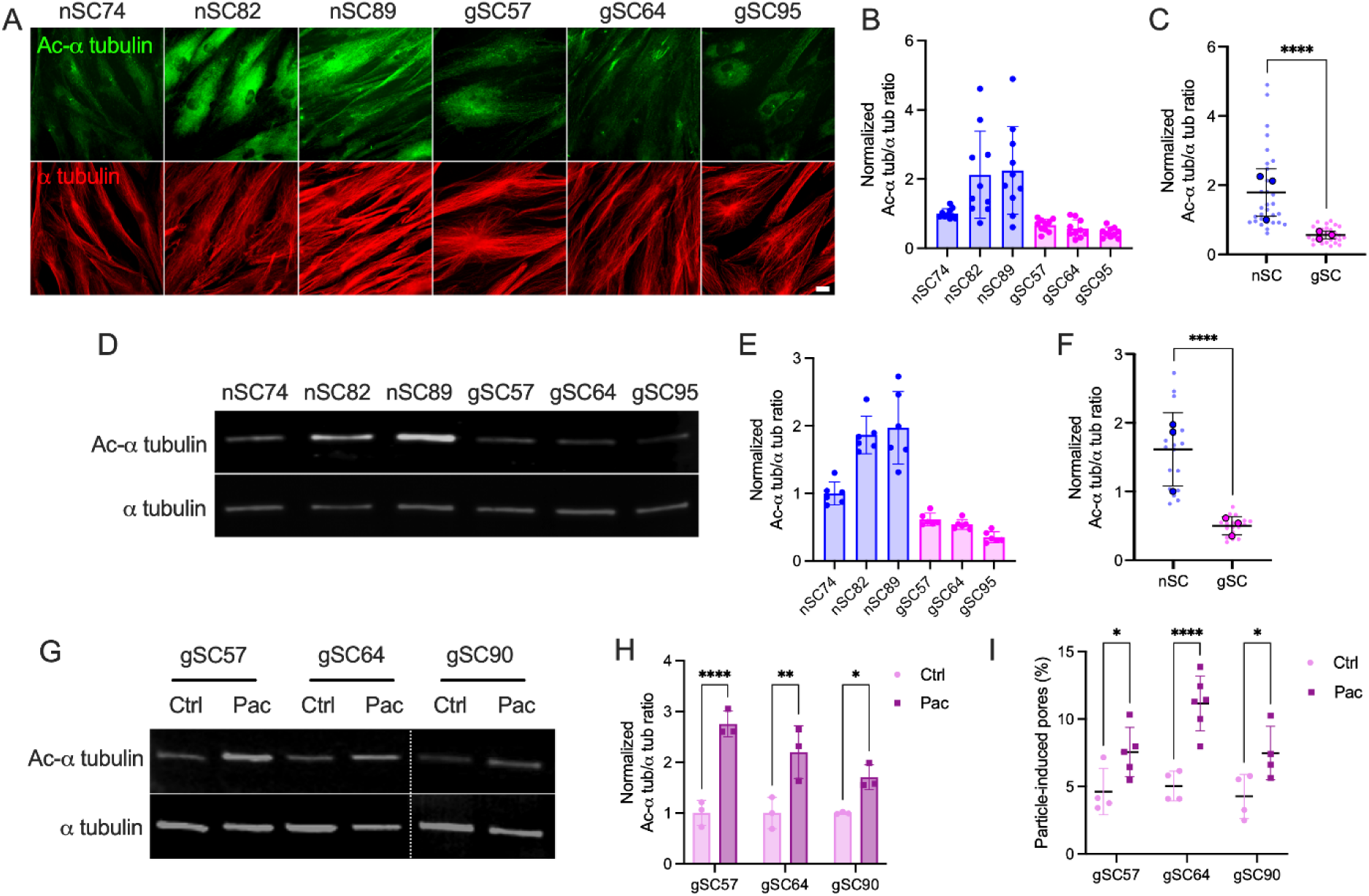
Glaucomatous SC (gSC) cells display reduced MT stability and increased pore formation rate upon MT stabilization. (A) Representative fluorescence micrographs of acetylated-α tubulin (ac-α tubulin) and total α tubulin in nSC and gSC cells. Scale bar, 20 μm. (B) Quantification of normalized ac-α tubulin/α tubulin fluorescence intensity (n = 10 images per group, with three independent replicates per strain). (C) Pooled analysis of normalized ac-α tubulin/α tubulin fluorescence intensity in nSC vs. gSC cells. Small dots represent technical replicates, and large dots represent the mean for each cell strain. (D) Immunoblots showing ac-α tubulin and total α tubulin in nSC and gSC cells. (E) Quantification of immunoblot data expressed as normalized ac-α tubulin/α tubulin ratios (n = 6 independent replicates). (F) Pooled analysis of normalized ac-α tubulin/α tubulin ratios from immunoblot experiments comparing nSC vs. gSC cells. Small dots represent technical replicates, and large dots represent the mean for each cell strain. (G) Representative immunoblot showing ac-α tubulin and total α tubulin in gSC cells with and without MT stabilization with 10 µM paclitaxel for 30 minutes. (H) Quantification of immunoblot data expressed as normalized ac-α tubulin/α tubulin ratios with and without MT stabilization (n = 3 independent replicates). (I) Pore incidence (ratio of pore-associated particles to particles under cells) in gSC cells with and without MT stabilization (n = 3-6 wells). Data are presented as mean ± SD. Statistical significance was determined using two-way ANOVA with multiple comparisons tests (*p<0.05, **p < 0.01, ****p < 0.0001).

We next evaluated the effects of the MT stabilizer paclitaxel in gSC cells. Paclitaxel treatment increased MT acetylation across the three gSC cell strains tested, indicating enhanced MT stability (Fig. 5G, H). Importantly, MT stabilization also significantly increased transcellular pore formation rate in gSC cells (Fig. 5I). Taken together, these findings demonstrate that SC cells from glaucoma patients exhibit reduced MT stability and that pharmacological MT stabilization enhances transcellular pore formation, suggesting a potential therapeutic avenue for restoring outflow function.

### MT stabilization increases ex vivo outflow facility in mouse eyes

Previous studies have shown that MT acetylation levels are inversely associated with trabecular meshwork cell contractility^53^. Consistent with this result, our findings demonstrate that MT stabilization decreases SC cell contractility and stiffness while enhancing transcellular pore formation. Based on these observations, we hypothesized that increasing MT stability would enhance AH outflow through the conventional pathway, consistent with previous findings showing that MT stabilization lowers IOP^53^. To test this hypothesis, we cannulated the anterior chambers of paired enucleated wild-type C57BL/6 mouse eyes and perfused them with either vehicle or paclitaxel for 45 minutes. MT stabilization significantly increased *ex vivo* outflow facility by 44.3% compared with vehicle-treated controls (Fig. 6A, B), indicating that enhancing MT stability improves conventional outflow function. We further confirmed that paclitaxel perfusion increased MT stabilization in the trabecular meshwork and SC inner wall region (Fig. 6C, D). Together, these findings suggest that paclitaxel enhances outflow facility in *ex vivo* mouse eyes by stabilizing MTs in conventional outflow tissues.

**Figure 6.**
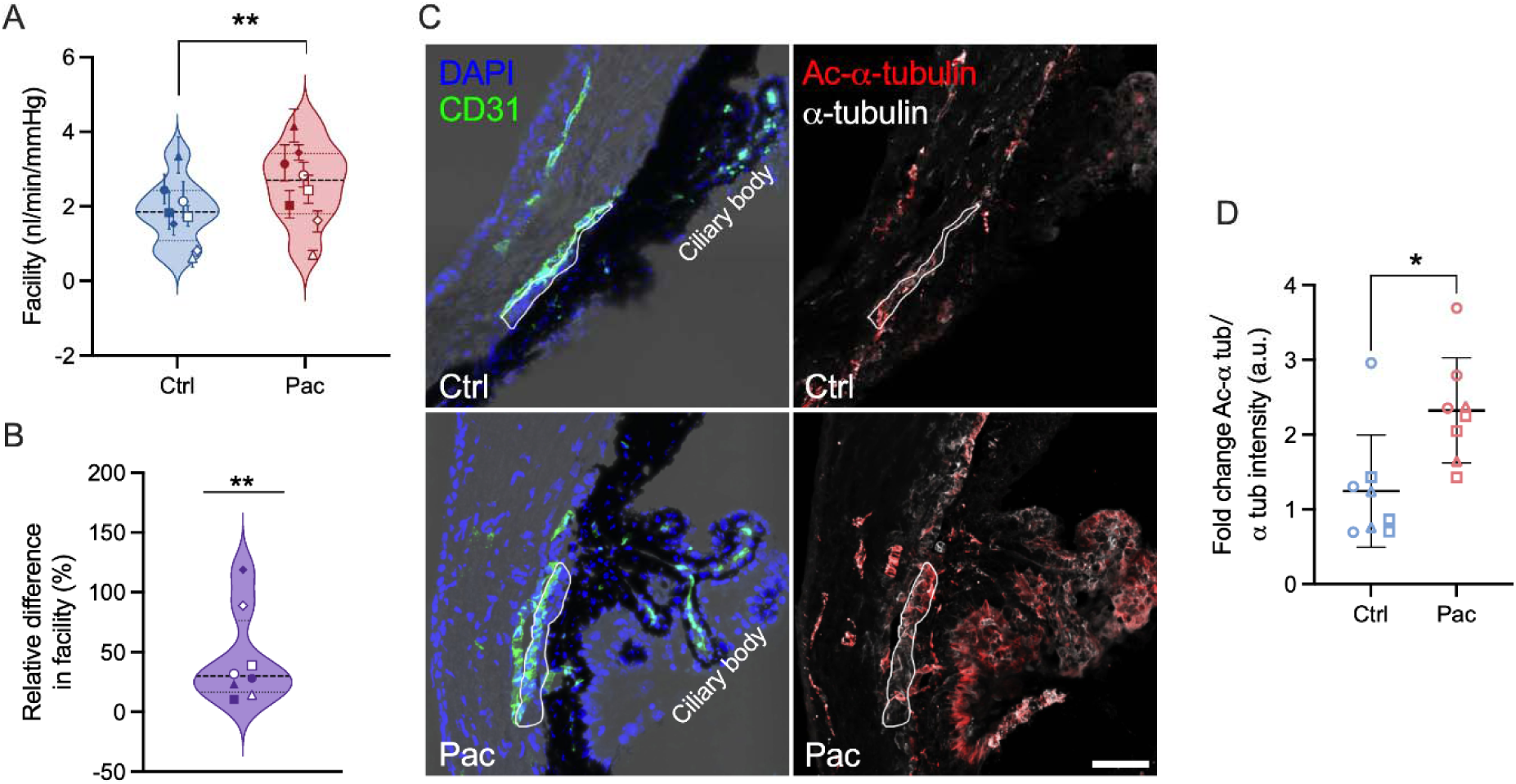
Perfusion with paclitaxel to enhance MT stabilization increases outflow facility in mouse eyes. (A) Outflow facility measured in C57BL/6J mouse eyes perfused with vehicle (1% DMSO) or 100 µM paclitaxel (N = 8 pairs of eyes). Outflow facility for each eye is shown as an individual data point with 95% confidence intervals. Different symbol shapes/fills indicate paired eyes. (B) Relative difference in outflow facility between paired eyes. Symbols correspond to the paired eyes shown in (A). Violin plots show the median and the 25th and 75th percentiles. (C) Representative immunofluorescence micrographs of frozen anterior segment sections stained for acetylated-α-tubulin (ac-α-tubulin; red) and total α-tubulin (pseudocolored grey to facilitate visualization of the red channel). CD31 staining (green) and brightfield imaging were used to visualize SC and identify the trabecular meshwork, and DAPI was used to label nuclei. The white line delineates the trabecular meshwork and SC inner wall. Scale bar, 50 μm. Note: In some sections, the SC lumen is not clearly visible, likely due to canal collapse during processing, causing contact between the SC inner and outer walls. CD31 was used to label SC endothelial cells, and in such cases, a line was drawn along the midpoint to delineate the inner wall. (D) Quantification of normalized ac-α tubulin/α tubulin fluorescence intensity in the trabecular meshwork and SC inner wall of perfused paired eyes (n = 2-3 sections per eye from three pairs of eyes). Different symbol shapes indicate paired eyes. Statistical significance was assessed using a paired t-test on log-transformed data in (A) and a nonparametric one-sample test against zero in (B), and two-way ANOVA with multiple comparisons in (D) (**p < 0.01, ****p < 0.0001).

## Discussion

In this study, we show that MT stability plays an important role in SC cell mechanobiology, transcellular pore formation, and outflow facility. Specifically, MT stabilization reduced actomyosin contractility and cellular stiffness in SC cells, whereas MT destabilization produced the opposite effects. These changes were mediated through the MT-associated Rho guanine nucleotide exchange factor GEF-H1. In SC cells derived from glaucoma patients, MT stability was reduced, suggesting that decreased MT stability may contribute to glaucoma pathology. Functionally, MT stabilization increased transcellular pore formation in SC cells from both healthy and glaucomatous donors and enhanced outflow facility in ex vivo mouse eyes. Together, these findings identify MT stability as an important determinant of SC inner wall biomechanics and AH outflow.

Both MTs and F-actin are key components of the cytoskeleton. MTs contribute to cell structure, intracellular transport, and spatial organization of signaling molecules, whereas F-actin generates contractile forces through actomyosin interactions^54^. Here, we show that MT stability influences F-actin stress fibers in SC cells. Long-term MT stabilization (i.e., 24 h) reduced F- actin polymerization, whereas MT destabilization increased actin stress fiber formation. Consistent with these changes, MT stabilization decreased p-MLC, while MT destabilization increased p-MLC levels. These results indicate that MT stability negatively modulates actomyosin contractility and thereby influences the mechanical phenotype of SC cells. Notably, SC cells derived from glaucoma patients exhibited reduced MT stability, suggesting potential increased cellular contractility consistent with the fibrotic phenotype previously reported in glaucomatous SC cells^52^. Conversely, ROCK signaling and F-actin dynamics may also influence MT stability^55–57^, a possibility that remains to be investigated in future studies.

Our results further identify GEF-H1 as a signaling link between MT dynamics and actomyosin contractility. GEF-H1 is a MT-associated RhoA exchange factor that becomes activated upon MT depolymerization^45,46^. We observed that GEF-H1 knockdown reduced p-MLC levels without altering F-actin levels. Although GEF-H1 activates RhoA upstream of both ROCK and formin-mediated actin polymerization pathways, this preferential effect may reflect differential sensitivity of downstream pathways to RhoA signaling in nSC cells. In addition, compensation by other RhoGEFs or parallel actin regulatory pathways may preserve F-actin levels, while the ROCK–myosin axis remains more dependent on GEF-H1 activity. Alternatively, GEF-H1 knockdown may alter actin organization rather than total actin abundance, while p-MLC may represent a more sensitive readout of reduced contractility than bulk F-actin intensity. Consistent with this mechanism, MT stabilizer-induced increases in actomyosin contractility were abolished by GEF-H1 knockdown. These findings suggest that MT destabilization activates RhoA-ROCK signaling through GEF-H1 release in SC cells. Similar MT-dependent regulation of GEF-H1 has been reported in other cell types, including fibroblasts^58^ and lymphocytes^59^, indicating that this pathway may represent a conserved mechanism linking MT dynamics to cellular contractility.

Primary cilia are microtubule-based sensory organelles that function as important mechanosensors in many cell types^60^. The ciliary axoneme is composed of highly acetylated microtubules, suggesting that MT stability may influence both the structure and function of primary cilia. In SC cells, primary cilia have been shown to play a role in shear stress-induced autophagy^61^, and loss of cilia in SC cells leads to decreased outflow facility^62^. These observations suggest that MT stability may influence SC cell mechanobiology not only through modulation of cytoskeletal mechanics but also through potential effects on cilia-dependent mechanosensory signaling.

The trabecular meshwork, which forms the substrate underlying SC inner wall cells, is markedly stiffer in glaucomatous eyes than in normal eyes^48–51^. To model this mechanical environment, we developed a biomimetic hydrogel system that mimics the stiffness of normal and glaucomatous trabecular meshwork^13^. Using this system, we previously demonstrated that normal SC cells cultured on substrates mimicking glaucomatous stiffness exhibit increased F-actin stress fiber formation and higher cellular stiffness^13^. Here, we further observed that normal SC cells cultured on stiff substrates displayed reduced MT stability, indicating that MTs in SC cells are mechanosensitive to substrate stiffness. Together with our findings linking MT stability to actomyosin contractility and cellular stiffness, these results suggest a potential bidirectional interplay between substrate stiffness and MT dynamics, in which substrate stiffness modulates MT stability, which in turn influences cytoskeletal organization and cell mechanics. Notably, MT responses to substrate stiffness appear to be cell type dependent. Similar to SC cells, epithelial cells have been reported to exhibit decreased MT stability when cultured on stiff substrates^26^. In contrast, fibroblasts^25^, B cells^63^ and astrocytes^24^ show increased MT stability under similar conditions, highlighting cell type-specific differences in MT-based mechano-responses.

SC inner wall cells facilitate AH drainage through the formation of pores, including both transcellular and paracellular pores^10^. Previous studies have shown that transcellular pore formation is strongly influenced by cellular stiffness and contractility^12,15^. In the present study, MT stabilization decreased cell contractility and softened SC cells. Consistent with these mechanical changes, MT stabilization increased transcellular pore formation in both normal and glaucomatous SC cells, whereas MT destabilization reduced pore incidence. These effects were consistent with MT stability-linked changes in cellular stiffness, in agreement with previous observations that softer SC cells exhibit increased pore formation. Because pores provide low-resistance pathways for AH flow across the SC inner wall, MT-dependent changes in cell mechanics are likely to influence outflow resistance.

In addition to modulating actomyosin mechanics, MT stability may influence transcellular pore formation through vesicle trafficking and membrane remodeling. Formation of transcellular pores likely requires membrane fusion of the basal and apical plasma membranes, or between the plasma membrane and intracellular vesicles. MTs serve as primary routes for intracellular vesicle transport, supporting kinesin- and dynein-mediated trafficking of membrane vesicles that help maintain plasma membrane homeostasis^64,65^. Stabilization of MTs may therefore facilitate transcellular pore formation by promoting vesicle delivery and membrane proteins redistribution during cellular deformation, whereas MT destabilization could impair vesicle transport and limit the membrane remodeling required for transcellular pore development. In addition, GEF-H1 has been reported to influence vesicle accumulation and organization in other cell types^66^. Further studies are needed to determine how MT stability modulates vesicle trafficking and membrane remodeling during transcellular pore formation.

Another important factor influencing transcellular pore formation in SC cells is intracellular Ca^2^. Ca^2^ has been shown to be essential for membrane fusion–driven transcellular formation and expansion^12,67^. Elevation of intracellular Ca^2^ promotes membrane fusion^68^, vesicle trafficking^69^, and cytoskeletal remodeling^70^, processes that are likely required for the formation and enlargement of transcellular pores spanning the basal and apical plasma membranes. In SC cells, mechanosensitive Ca^2^ influx has been shown to facilitate transcellular pore formation^13^. Given the known interactions between microtubules and mechanosensitive signaling pathways, MT stability may also influence Ca^2^ -dependent processes during transcellular pore formation. For example, MTs may modulate the trafficking and spatial organization of ion channels and signaling complexes, thereby modulating mechanosensitive Ca^2^ influx. How MT dynamics interact with Ca^2^ signaling during transcellular pore formation in SC cells remains to be determined.

Finally, MT stabilization increased outflow facility in ex vivo mouse eyes. Paclitaxel increased facility relative to contralateral control eyes, indicating that MT-dependent cytoskeletal remodeling can influence AH outflow at the tissue level. Because perfused paclitaxel also reaches the trabecular meshwork, the increase in facility may not arise solely from effects on the SC inner wall but may also involve trabecular meshwork responses. Consistent with this interpretation, previous studies have shown that MT stability affects trabecular meshwork cell contractility and that MT stabilization can lower IOP in mouse eyes^53^. Although paclitaxel itself may not be suitable for therapeutic use, as it can be cytotoxic and interfere with cell division in ocular tissues such as corneal endothelial and trabecular meshwork cells^71,72^, these findings suggest that MT-related pathways may represent potential targets for improving outflow function. In particular, modulators such as GEF-H1 that influence MT stability may provide a more selective strategy to modulate SC cell mechanics and outflow.

SC inner wall cells facilitate AH drainage through pressure-dependent pore formation. Altered cellular biomechanics in these cells contributes to increased outflow resistance and elevated IOP in glaucoma. This study identifies MT stability as a modulator of SC cell mechanics, linking MT dynamics to actomyosin contractility, cellular stiffness, and transcellular pore formation. Pharmacological stabilization of MTs increased transcellular pore formation and enhanced outflow facility in mouse eyes. These findings reveal a previously unrecognized cytoskeletal mechanism modulating AH outflow and suggest that MT-dependent signaling pathways may represent potential targets for improving outflow function in glaucoma.

## Disclosure

The authors report no conflicts of interest.

## Acknowledgments

This study was financially supported by BrightFocus Foundation G2024003F (HL); National Institutes of Health R01EY028608 (WDS), R01EY022359 (WDS), R01EY030124 (CRE and WDS), R21EY035468 (AJF), and R01EY030871 (AJF); Vision Training Grant T32 EY007092- 38 (NSFG) and the Alfred P. Sloan Foundation G-2019-11435 (NSFG); the National Institutes of Health NEI P30 core grant P30EY006360, and a Challenge Grant from Research to Prevent Blindness, Inc. to the Department of Ophthalmology at Emory University, Georgia Research Alliance (CRE), and Emory Ophthalmology Departmental funds (CRE).

## Author contributions

**Conceptualization:** H.L., C.R.E.; **Investigation:** H.L., N.S.F.-G., K.M.P. M.C., W.D.S., C.R.E. designed all experiments, collected, analyzed, and interpreted the data. N.S.F.-G. assisted with the outflow facility measurements using the iPerfusion system and with data analysis. K.M.P. isolated and characterized the primary human SC cells. M.C. performed confocal imaging for the mouse eye anterior sections following outflow facility measurements; **Writing – original draft:** H.L., C.R.E.; **Writing – review and editing:** all authors. **Supervision:** C.R.E.

## Notes

### Competing Interest Statement

The authors have declared no competing interest.

## References

1. Quigley HA, Broman AT. The number of people with glaucoma worldwide in 2010 and 2020. Br J Ophthalmol. Mar 2006;90(3):262–267. doi:10.1136/bjo.2005.081224

2. Braunger BM, Fuchshofer R, Tamm ER. The aqueous humor outflow pathways in glaucoma: A unifying concept of disease mechanisms and causative treatment. European Journal of Pharmaceutics and Biopharmaceutics. 2015/09/01/ 2015;95:173–181. 10.1016/j.ejpb.2015.04.029

3. Stamer WD. The Cell and Molecular Biology of Glaucoma: Mechanisms in the Conventional Outflow Pathway. Investigative Ophthalmology & Visual Science. 2012;53(5):2470–2472. doi:10.1167/iovs.12-9483f

4. Ashpole NE, Overby DR, Ethier CR, Stamer WD. Shear stress-triggered nitric oxide release from Schlemm’s canal cells. Investigative ophthalmology & visual science. 2014;55(12):8067–8076. doi:10.1167/iovs.14-14722

5. Dautriche CN, Tian Y, Xie Y, Sharfstein ST. A Closer Look at Schlemm’s Canal Cell Physiology: Implications for Biomimetics. J Funct Biomater. Sep 21 2015;6(3):963–985. doi:10.3390/jfb6030963

6. Stamer WD, Braakman ST, Zhou EH, et al. Biomechanics of Schlemm’s canal endothelium and intraocular pressure reduction. Prog Retin Eye Res. Jan 2015;44:86–98. doi:10.1016/j.preteyeres.2014.08.002

7. Ethier CR. The inner wall of Schlemm’s canal. Exp Eye Res. Feb 2002;74(2):161–172. doi:10.1006/exer.2002.1144

8. Stamer WD, Roberts BC, Howell DN, Epstein DL. Isolation, culture, and characterization of endothelial cells from Schlemm’s canal. Invest Ophthalmol Vis Sci. Sep 1998;39(10):1804–1812.

9. Ethier CR, Coloma FM, Sit AJ, Johnson M. Two pore types in the inner-wall endothelium of Schlemm’s canal. Invest Ophthalmol Vis Sci. Oct 1998;39(11):2041–2048.

10. Holmberg A. The Fine Structure of the Inner Wall of Schlemm’s Canal. AMA Archives of Ophthalmology. 1959;62(6):956–958. doi:10.1001/archopht.1959.04220060028005

11. Johnson M, Chan D, Read AT, Christensen C, Sit A, Ethier CR. The Pore Density in the Inner Wall Endothelium of Schlemm’s Canal of Glaucomatous Eyes. Investigative Ophthalmology & Visual Science. 2002;43(9):2950–2955.

12. Siadat SM, Li H, Millette B, et al. Endothelial cell stiffness and type drive the formation of biomechanically induced transcellular pores. Cell Reports. 2026;45(1)doi:10.1016/j.celrep.2025.116668

13. Li H, Wong C, Siadat SM, et al. Transient receptor potential vanilloid 4 modulates substrate stiffness mechanosensing and transcellular pore formation in human Schlemm’s canal cells. Acta Biomaterialia. 2025/09/27/ 2025;10.1016/j.actbio.2025.09.039

14. Zhou EH, Krishnan R, Stamer WD, et al. Mechanical responsiveness of the endothelial cell of Schlemm’s canal: scope, variability and its potential role in controlling aqueous humour outflow. J R Soc Interface. Jun 7 2012;9(71):1144–1155. doi:10.1098/rsif.2011.0733

15. Overby DR, Zhou EH, Vargas-Pinto R, et al. Altered mechanobiology of Schlemm’s canal endothelial cells in glaucoma. Proc Natl Acad Sci U S A. Sep 23 2014;111(38):13876–13881. doi:10.1073/pnas.1410602111

16. Gardner MK, Zanic M, Howard J. Microtubule catastrophe and rescue. Current Opinion in Cell Biology. 2013/02/01/ 2013;25(1):14–22. 10.1016/j.ceb.2012.09.006

17. Hawkins T, Mirigian M, Selcuk Yasar M, Ross JL. Mechanics of microtubules. Journal of Biomechanics. 2010/01/05/ 2010;43(1):23–30. 10.1016/j.jbiomech.2009.09.005

18. Mitchison T, Kirschner M. Dynamic instability of microtubule growth. Nature. 1984/11/01 1984;312(5991):237–242. doi:10.1038/312237a0

19. Fletcher DA, Mullins RD. Cell mechanics and the cytoskeleton. Nature. 2010/01/01 2010;463(7280):485–492. doi:10.1038/nature08908

20. Fourriere L, Jimenez AJ, Perez F, Boncompain G. The role of microtubules in secretory protein transport. J Cell Sci. Jan 29 2020;133(2)doi:10.1242/jcs.237016

21. Garcin C, Straube A. Microtubules in cell migration. Essays Biochem. Oct 31 2019;63(5):509–520. doi:10.1042/ebc20190016

22. Bouchet BP, Akhmanova A. Microtubules in 3D cell motility. J Cell Sci. Jan 1 2017;130(1):39–50. doi:10.1242/jcs.189431

23. Lyons JS, Joca HC, Law RA, et al. Microtubules tune mechanotransduction through NOX2 and TRPV4 to decrease sclerostin abundance in osteocytes. Sci Signal. Nov 21 2017;10(506)doi:10.1126/scisignal.aan5748

24. Seetharaman S, Vianay B, Roca V, et al. Microtubules tune mechanosensitive cell responses. Nat Mater. Mar 2022;21(3):366–377. doi:10.1038/s41563-021-01108-x

25. Wen D, Gao Y, Liu Y, et al. Matrix stiffness-induced α-tubulin acetylation is required for skin fibrosis formation through activation of Yes-associated protein. MedComm (2020). Aug 2023;4(4):e319. doi:10.1002/mco2.319

26. Heck JN, Ponik SM, Garcia-Mendoza MG, et al. Microtubules regulate GEF-H1 in response to extracellular matrix stiffness. Mol Biol Cell. Jul 2012;23(13):2583–2592. doi:10.1091/mbc.E11-10-0876

27. Coleman AK, Joca HC, Shi G, Lederer WJ, Ward CW. Tubulin acetylation increases cytoskeletal stiffness to regulate mechanotransduction in striated muscle. J Gen Physiol. Jul 5 2021;153(7)doi:10.1085/jgp.202012743

28. Li H, Singh A, Perkumas KM, Stamer WD, Ganapathy PS, Herberg S. YAP/TAZ Mediate TGFβ2-Induced Schlemm’s Canal Cell Dysfunction. Investigative Ophthalmology & Visual Science. 2022;63(12):15–15. doi:10.1167/iovs.63.12.15

29. Li H, Henty-Ridilla JL, Bernstein AM, Ganapathy PS, Herberg S. TGFβ2 regulates human trabecular meshwork cell contractility via ERK and ROCK pathways with distinct signaling crosstalk dependent on the culture substrate. Current Eye Research. 2022:1–41. doi:10.1080/02713683.2022.2071943

30. Li H, Raghunathan V, Stamer WD, Ganapathy PS, Herberg S. Extracellular Matrix Stiffness and TGFβ2 Regulate YAP/TAZ Activity in Human Trabecular Meshwork Cells. Original Research. Frontiers in Cell and Developmental Biology. 2022-March-01 2022;10doi:10.3389/fcell.2022.844342

31. Li H, Kuhn M, Kelly RA, et al. Targeting YAP/TAZ mechanosignaling to ameliorate stiffness-induced Schlemm’s canal cell pathobiology. American Journal of Physiology-Cell Physiology. 2024;326(2):C513–C528. doi:10.1152/ajpcell.00438.2023

32. Singh A, Ghosh R, Li H, et al. Three-Dimensional Extracellular Matrix Protein Hydrogels for Human Trabecular Meshwork Cell Studies. In: Jakobs T, Liton PB, eds. Glaucoma: Methods and Protocols. New York, NY: Springer US; 2025:17-29.

33. Li H, Harvey DH, Dai J, et al. Characterization, Enrichment, and Computational Modeling of Cross-Linked Actin Networks in Transformed Trabecular Meshwork Cells. Investigative Ophthalmology & Visual Science. 2025;66(2):65–65. doi:10.1167/iovs.66.2.65

34. Schindelin J, Arganda-Carreras I, Frise E, et al. Fiji: an open-source platform for biological-image analysis. Nat Methods. Jun 28 2012;9(7):676–682. doi:10.1038/nmeth.2019

35. Vargas-Pinto R, Gong H, Vahabikashi A, Johnson M. The effect of the endothelial cell cortex on atomic force microscopy measurements. Biophys J. Jul 16 2013;105(2):300–309. doi:10.1016/j.bpj.2013.05.034

36. Siadat SM, Li H, Millette BA, et al. Endothelial cell stiffness and type drive the formation of biomechanically-induced transcellular pores. bioRxiv. 2024:2024.2010.2023.619950. doi:10.1101/2024.10.23.619950

37. Sherwood JM, Reina-Torres E, Bertrand JA, Rowe B, Overby DR. Measurement of outflow facility using iPerfusion. PLoS One. 2016;11(3):e0150694.

38. Kim D, Fang R, Zhang P, et al. In Vivo Quantification of Anterior and Posterior Chamber Volumes in Mice: Implications for Aqueous Humor Dynamics. Investigative Ophthalmology & Visual Science. 2025;66(1):18–18. doi:10.1167/iovs.66.1.18

39. Wong CA, Fraticelli Guzmán NS, Read AT, et al. A method for analyzing AFM force mapping data obtained from soft tissue cryosections. Journal of Biomechanics. 2024/05/01/ 2024;168:112113. 10.1016/j.jbiomech.2024.112113

40. Eshun-Wilson L, Zhang R, Portran D, et al. Effects of α-tubulin acetylation on microtubule structure and stability. Proceedings of the National Academy of Sciences. 2019;116(21):10366–10371. doi:doi:10.1073/pnas.1900441116

41. Saunders HAJ, Johnson-Schlitz DM, Jenkins BV, Volkert PJ, Yang SZ, Wildonger J. Acetylated α-tubulin K394 regulates microtubule stability to shape the growth of axon terminals. Curr Biol. Feb 7 2022;32(3):614–630.e615. doi:10.1016/j.cub.2021.12.012

42. Kühn S, Geyer M. Formins as effector proteins of Rho GTPases. Small GTPases. 2014;5:e29513. doi:10.4161/sgtp.29513

43. Kümper S, Mardakheh FK, McCarthy A, et al. Rho-associated kinase (ROCK) function is essential for cell cycle progression, senescence and tumorigenesis. eLife. 2016/01/14 2016;5:e12203. doi:10.7554/eLife.12203

44. Lawson CD, Ridley AJ. Rho GTPase signaling complexes in cell migration and invasion. J Cell Biol. Feb 5 2018;217(2):447–457. doi:10.1083/jcb.201612069

45. Birkenfeld J, Nalbant P, Yoon SH, Bokoch GM. Cellular functions of GEF-H1, a microtubule-regulated Rho-GEF: is altered GEF-H1 activity a crucial determinant of disease pathogenesis? Trends Cell Biol. May 2008;18(5):210–219. doi:10.1016/j.tcb.2008.02.006

46. Meiri D, Marshall CB, Mokady D, et al. Mechanistic insight into GPCR-mediated activation of the microtubule-associated RhoA exchange factor GEF-H1. Nat Commun. Sep 11 2014;5:4857. doi:10.1038/ncomms5857

47. Johnson M, Chan D, Read AT, Christensen C, Sit A, Ethier CR. The pore density in the inner wall endothelium of Schlemm’s canal of glaucomatous eyes. Invest Ophthalmol Vis Sci. Sep 2002;43(9):2950–2955.

48. Vahabikashi A, Gelman A, Dong B, et al. Increased stiffness and flow resistance of the inner wall of Schlemm’s canal in glaucomatous human eyes. Proceedings of the National Academy of Sciences. 2019;116(52):26555–26563. doi:10.1073/pnas.1911837116

49. Wang K, Johnstone MA, Xin C, et al. Estimating Human Trabecular Meshwork Stiffness by Numerical Modeling and Advanced OCT Imaging. Investigative Ophthalmology & Visual Science. 2017;58(11):4809–4817. doi:10.1167/iovs.17-22175

50. Wang K, Read AT, Sulchek T, Ethier CR. Trabecular meshwork stiffness in glaucoma. Exp Eye Res. May 2017;158:3–12. doi:10.1016/j.exer.2016.07.011

51. Last JA, Pan T, Ding Y, et al. Elastic Modulus Determination of Normal and Glaucomatous Human Trabecular Meshwork. Investigative Ophthalmology & Visual Science. 2011;52(5):2147–2152. doi:10.1167/iovs.10-6342

52. Kelly RA, Perkumas KM, Campbell M, et al. Fibrotic Changes to Schlemm’s Canal Endothelial Cells in Glaucoma. Int J Mol Sci. Aug 31 2021;22(17)doi:10.3390/ijms22179446

53. Su CC, Desikan V, Betsch K, Shim MS, Keller KE, Liton PB. Tubulin Acetylation Enhances Microtubule Stability in Trabecular Meshwork Cells Under Mechanical Stress. Invest Ophthalmol Vis Sci. Jan 2 2025;66(1):43. doi:10.1167/iovs.66.1.43

54. Goode BL, Drubin DG, Barnes G. Functional cooperation between the microtubule and actin cytoskeletons. Current Opinion in Cell Biology. 2000/02/01/ 2000;12(1):63–71. 10.1016/S0955-0674(99)00058-7

55. Kadir S, Astin JW, Tahtamouni L, Martin P, Nobes CD. Microtubule remodelling is required for the front-rear polarity switch during contact inhibition of locomotion. J Cell Sci. Aug 1 2011;124(Pt 15):2642–2653. doi:10.1242/jcs.087965

56. Xu Z, Schaedel L, Portran D, et al. Microtubules acquire resistance from mechanical breakage through intralumenal acetylation. Science. Apr 21 2017;356(6335):328–332. doi:10.1126/science.aai8764

57. Scaife RM, Job D, Langdon WY. Rapid Microtubule-dependent Induction of Neurite-like Extensions in NIH 3T3 Fibroblasts by Inhibition of ROCK and Cbl. Mol Biol Cell. 2003;14(11):4605–4617. doi:10.1091/mbc.e02-11-0739

58. Chang YC, Nalbant P, Birkenfeld J, Chang ZF, Bokoch GM. GEF-H1 couples nocodazole-induced microtubule disassembly to cell contractility via RhoA. Mol Biol Cell. May 2008;19(5):2147–2153. doi:10.1091/mbc.e07-12-1269

59. Pineau J, Pinon L, Mesdjian O, Fattaccioli J, Lennon Duménil A-M, Pierobon P. Microtubules restrict F-actin polymerization to the immune synapse via GEF-H1 to maintain polarity in lymphocytes. eLife. 2022/09/16 2022;11:e78330. doi:10.7554/eLife.78330

60. Kiesel P, Alvarez Viar G, Tsoy N, et al. The molecular structure of mammalian primary cilia revealed by cryo-electron tomography. Nat Struct Mol Biol. Dec 2020;27(12):1115–1124. doi:10.1038/s41594-020-0507-4

61. Shim MS, Dixon A, Nettesheim A, et al. Shear stress induces autophagy in Schlemm’s canal cells via primary cilia-mediated SMAD2/3 signaling pathway. Autophagy Rep. 2023;2(1)doi:10.1080/27694127.2023.2236519

62. Kizhatil K, Clark G. Primary cilia and aqueous humor outflow. Investigative Ophthalmology & Visual Science. 2024;65(7):4261–4261.

63. Aceitón P, Riobó I, Del Valle Batalla F, et al. B cell mechanotransduction via ATAT1 coordinates actin and lysosomal dynamics at the immune synapse. J Cell Biol. Aug 4 2025;224(8)doi:10.1083/jcb.202407181

64. Verdeny-Vilanova I, Wehnekamp F, Mohan N, et al. 3D motion of vesicles along microtubules helps them to circumvent obstacles in cells. J Cell Sci. Jun 1 2017;130(11):1904–1916. doi:10.1242/jcs.201178

65. D’Souza AI, Grover R, Monzon GA, Santen L, Diez S. Vesicles driven by dynein and kinesin exhibit directional reversals without regulators. Nature Communications. 2023/11/20 2023;14(1):7532. doi:10.1038/s41467-023-42605-8

66. Pathak R, Delorme-Walker VD, Howell MC, et al. The microtubule-associated Rho activating factor GEF-H1 interacts with exocyst complex to regulate vesicle traffic. Dev Cell. Aug 14 2012;23(2):397–411. doi:10.1016/j.devcel.2012.06.014

67. Lai Y, Diao J, Liu Y, et al. Fusion pore formation and expansion induced by Ca^2+^ and synaptotagmin 1. Proceedings of the National Academy of Sciences. 2013;110(4):1333–1338. doi:doi:10.1073/pnas.1218818110

68. Tahir MA, Guven ZP, Arriaga LR, et al. Calcium-triggered fusion of lipid membranes is enabled by amphiphilic nanoparticles. Proceedings of the National Academy of Sciences. 2020;117(31):18470–18476. doi:doi:10.1073/pnas.1902597117

69. Sargeant J, Hay JC. Ca(2+) regulation of constitutive vesicle trafficking. Fac Rev. 2022;11:6. doi:10.12703/r/11-6

70. Tsai F-C, Kuo G-H, Chang S-W, Tsai P-J. Ca2+ Signaling in Cytoskeletal Reorganization, Cell Migration, and Cancer Metastasis. BioMed Research International. 2015/01/01 2015;2015(1):409245. 10.1155/2015/409245

71. Choritz L, Grub J, Wegner M, Pfeiffer N, Thieme H. Paclitaxel inhibits growth, migration and collagen production of human Tenon’s fibroblasts--potential use in drug- eluting glaucoma drainage devices. Graefes Arch Clin Exp Ophthalmol. Feb 2010;248(2):197–206. doi:10.1007/s00417-009-1221-4

72. Pasquier E, Carré M, Pourroy B, et al. Antiangiogenic activity of paclitaxel is associated with its cytostatic effect, mediated by the initiation but not completion of a mitochondrial apoptotic signaling pathway. Mol Cancer Ther. Oct 2004;3(10):1301–1310.

