## Supplemental file for "Microtubule stability modulates Schlemm’s canal cell mechanobiology and outflow facility in glaucoma"

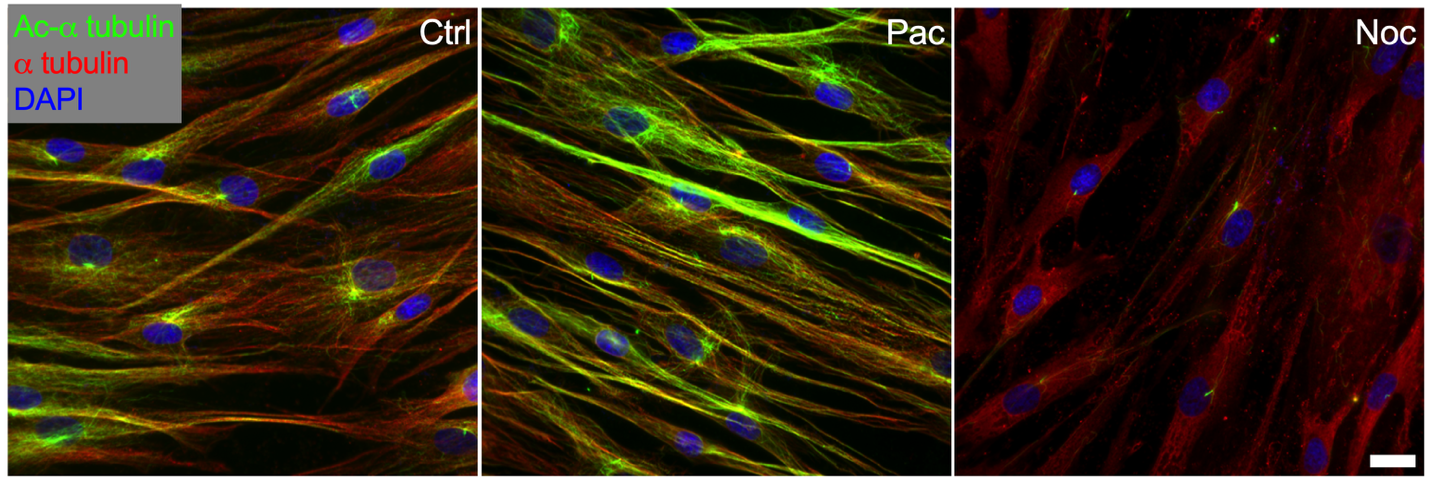

**Suppl. Fig. 1. Paclitaxel and nocodazole affect tubulin acetylation levels in nSC cells.** Representative fluorescence micrographs of acetylated-α tubulin (ac-α tubulin) and total α tubulin in normal SC (nSC) cells following 30-minute treatment with control (DMSO), 10 μM paclitaxel (Pac), or 10 μM nocodazole (Noc). Scale bar, 20 μm.

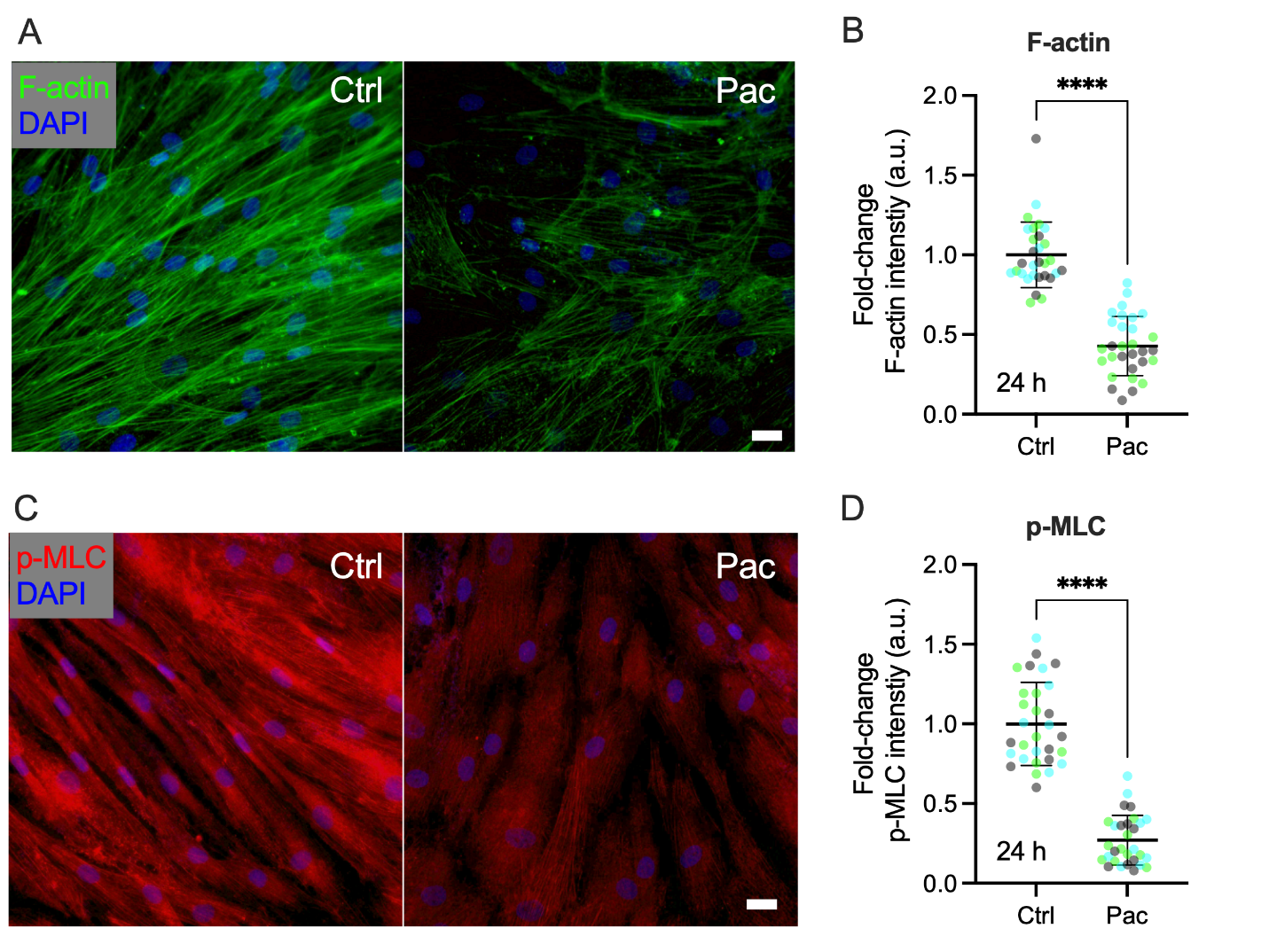

**Suppl. Fig. 2. Prolonged MT stabilization exerts a stronger effect on F-actin and p-MLC levels than acute stabilization.** (A, C) Representative fluorescence micrographs of F-actin and phosphorylated myosin light chain (p-MLC) in normal SC (nSC) cells treated with control (DMSO) or 10 μM paclitaxel (Pac) for 24 hours. Scale bar, 25 μm. (B, D) Quantification of normalized F-actin and p-MLC fluorescence intensities (n = 30 images per group from 3 different nSC cell strains, with three independent replicates per strain). Symbols of the same color represent data from the same cell strain. Data are presented as mean ± SD. Statistical significance was determined by two-way ANOVA with multiple comparisons tests (****p < 0.0001).
